# Placental treatment with *insulin-like growth factor 1* via nanoparticle differentially impacts vascular remodeling factors in guinea pig sub-placenta/decidua

**DOI:** 10.1101/2022.09.26.509492

**Authors:** Baylea Davenport, Helen N. Jones, Rebecca L. Wilson

## Abstract

Clinically, fetal growth restriction (FGR) is only detectable in later gestation, despite pathophysiological establishment likely earlier in pregnancy. Additionally, there are no effective in utero treatment options for FGR. We have developed a nanoparticle to deliver *human insulin-like 1* growth *factor* (*hIGF-1*) in a trophoblast-specific manner which results in increased expression of *hIGF-1*. IGF-1 signaling in the placenta regulates multiple developmental processes including trophoblast invasion and maternal vascular remodeling, both of which can be diminished in the FGR placenta. We aimed to determine the effects of short-term *hIGF-1* nanoparticle treatment on sub-placenta/decidua trophoblast signaling mechanisms in FGR and under normal growth conditions. Using the guinea pig maternal nutrient restriction (MNR) model of FGR, ultrasound-guided, intra-placenta injections of *hIGF-1* nanoparticle were performed at gestational day 30-33, and dams sacrificed five days later. Sub-placenta/decidua tissue was separated from placenta for further analyses. Western blot was used to analyze protein expression of ERK/AKT/mTOR signaling proteins (phospho-Erk (pErk), phospho-Akt (pAkt), raptor, rictor and deptor). qPCR was used to analyze gene expression of vascular/remodeling factors (*vascular endothelial growth factor* (*Vegf*), *placenta growth factor* (*Pgf*), *platelet-derived growth factor* (*Pdgf*)) and tight junction/adhesion proteins (*claudin 5* (*Cldn5*), *p-glycoprotein* (*Abcb1*), *occludin* (*Ocln*) and *tight junction protein 1* (*Zo1*)). MNR reduced expression of pErk, *PdgfB* and *Cldn5*, and increased expression of *Ocln* and *Zo1* in the sub-placenta/decidua. In MNR + *hIGF1* nanoparticle sub-placenta/decidua, expression of *PdgfB, Ocln* and *Zo1* was normalized, whilst pAkt, *VegfB, Vegf receptor 1* and *PdgfB receptor* were increased compared to MNR. In contrast, *hIGF-1* nanoparticle treatment of normal placentas reduced expression of pErk, raptor and increased expression of the mTOR inhibitor deptor. This was associated with reduced expression of *VegfA, Plgf*, and *PdgfB*. Here we have shown that the impact of *hIGF-1* nanoparticle treatment is dependent on pregnancy environment. Under MNR/FGR, *hIGF-1* nanoparticle treatment triggers increased expression of growth factors and normalization of EMT factors. However, under normal conditions, the response of the placenta is to decrease AKT/mTOR signaling and growth factor expression to achieve homeostasis.

## Introduction

Fetal growth restriction (FGR; estimated fetal weight <10th percentile), is a significant risk factor for stillbirth and contributes significantly to infant morbidity and mortality (Bernstein, Horbar et al. 2000). Currently, there is no effective in utero treatment for FGR, which in part, is due to the fact that whilst the pathophysiology of FGR is established in the first half of pregnancy, clinically FGR is only detectable in later gestation (Sun, Groom et al. 2020).

Fetal growth increases exponentially throughout the latter half of pregnancy (Reynolds and Redmer 1995), and increased uterine blood flow is essential to meet metabolic demands from the growing fetus, as well as uterus and placenta (Bulmer, Burton et al. 2012). Increased uterine blood flow is supported in part in humans, by properly remodeled maternal spiral arteries coordinated by invasion of extravillous trophoblasts (EVTs) (Burton and Fowden 2015). Aberrations in invasion can limit remodeling and hence, blood flow from the mother to the maternal-fetal interface, restricting nutrient and oxygen delivery and contributing to FGR (Brosens, Dixon et al. 1977, Kaufmann, Black et al. 2003). The guinea pig maternal nutrient restriction (MNR) model is an established model to study the effects of FGR on fetal development, pregnancy outcome and the developmental origins of health and disease (Kunkele and Trillmich 1997, Morrison, Botting et al. 2018). Moderate food restriction prior to and throughout pregnancy results in reduced fetal, and placental weight from mid gestation (Sohlstrom, Katsman et al. 1998) without significantly affecting fertility (Elias, Ghaly et al. 2016). Most importantly, placental development in the guinea pig includes trophoblast invasion, which is not present in other small animal models (Mess 2007). Defining the guinea pig placenta is the sub-placenta (Carter, Enders et al. 2006). This region, distinct from the chorioallantoic placenta, is comprised of trophoblast arranged in folded layers encircled by syncytial trophoblast, and the source of invasive trophoblasts (Verkeste, Slangen et al. 1998). These invasive trophoblasts are the source of interstitial invasion into the decidua, and into the arterial walls similar to human placentation (Carter, Enders et al. 2006), making guinea pigs ideal for identifying mechanisms involved in trophoblast invasion.

One of the central signaling cascades that regulates trophoblast invasion is the ERK/AKT signaling pathway (Knöfler and Pollheimer 2012). ERK/AKT signaling activates extracellular matrix breakdown, adhesion molecule reduction for epithelial-mesenchymal transition (EMT) and invasion, and mTOR/raptor/rictor activation for proliferation (Chakraborty, Gleeson et al. 2002, Xu, Yang et al. 2015, Crespo*, Kind* et al. 2016). Each of these pathways are critical for EVTs to invade into the maternal tissue. Moreover, trophoblast invasion can be intrinsically and extrinsically influenced by growth factors, adhesion and tight junction molecules as well as extracellular matrix components (Lunghi, Ferretti et al. 2007). Most studied are the vascular endothelial growth factor (VEGF) proteins which include placental growth factor (PGF), and regulate placental development through binding with their specific transmembrane receptors, including VEGFR-1 (Flt-1) and VEGFR-2 (KDR) (Shore, Wang et al. 1997).

Tight junction and adhesion proteins are critical components in determining epithelial cell polarity. These tight junction proteins must be repressed for cells to go through EMT to reduce cell adhesion and increase cell mobility (migration/invasion) (Ikenouchi, Matsuda et al. 2003). Claudin protein expression has been shown to increase cell proliferation, invasion, and migration of EVTs in vitro. Through activation of the ERK1/2 pathway, claudin proteins upregulate matrix-metalloproteinases to help break down the extracellular matrix (ECM) as they invade into the maternal tissue and control permeability. mRNA levels of these claudin proteins have also been shown to be down in cases of hypertension and/or preeclampsia, suggesting a role in placental dysfunction (Zhao, Qi et al. 2020). During normal placenta development ZO-1 and occludin are expressed in the proliferating trophoblasts and down regulation of these proteins is associated with the epithelial cell transformation that is necessary for proper invasion of EVTs (Marzioni, Banita et al. 2001). P-glycoprotein is also down regulated in cases of preeclampsia and functionally important for EVT invasion (Ceckova-Novotna, Pavek et al. 2006, Dunk, Pappas et al. 2018).

Insulin-like 1 growth factor (IGF-1) is a known activator of the ERK/AKT signaling cascade in the maternal fetal interface (Dumolt, Powell et al. 2021). Expressed in placental and fetal tissue from the pre-implantation period through to birth (Fowden 2003), IGF-1 is known to regulate trophoblast invasion through binding with the IGF-1 receptor (IGF-1R). Fetal growth positively correlates with maternal IGF-1 concentrations (Chellakooty, Vangsgaard et al. 2004), and both maternal serum levels of IGF-1 and placenta expression of IGF-1R are decreased in FGR (Holmes, Montemagno et al. 1997, Laviola, Perrini et al. 2005). Additionally, we have previously shown that treatment of the placenta with a non-viral, polymer based, nanoparticle gene therapy that specifically increases *human IGF-1* (*hIGF-1*) in trophoblast, maintains fetal growth in a surgically-ligated FGR mouse model, and increases nutrient transporter expression (Abd Ellah, Taylor et al. 2015, Wilson, Owens et al. 2020, Wilson, Lampe et al. 2021). Placenta treatment with the *hIGF-1* nanoparticle does not adversely impact maternal or fetal outcomes, and cannot cross the placenta into fetal circulation (Wilson, Lampe et al. 2021, Wilson, Stephens et al. 2021). We have previously demonstrated that short-term treatment with the *hIGF-1* nanoparticle results in increased hIGF-1 expression in trophoblasts of the guinea pig sub-placenta/decidua (Wilson, Lampe et al. 2021). In the present study, we determined the effects of short-term *hIGF-1* nanoparticle treatment on trophoblast ERK/AKT signaling, and downstream effectors that regulate trophoblast invasion in both normal and FGR conditions.

## Methods

### Animals

All experiments were approved by the Cincinnati Children’s Hospital Medical Center and University of Florida Institutional Animal Care and Use Committee (Protocol numbers 2017-0065 and 202011236, respectively). Female Dunkin-Hartley guinea pigs were purchased (Charles River Laboratories, Wilmington, MA) at 500-550 g and housed in an environmentally controlled room (22°C/72°F and 50% humidity) under a 12 h light-dark cycle. Upon arrival, food (LabDiet diet 5025: 27% protein, 13.5% fat, and 60% carbohydrate as % of energy) and water were provided *ad libitum*. After a 2-week acclimatization period, females were weighed and assigned to either an *ad libitum* fed diet (termed Control: *n* = 11) or maternal nutrient restricted (MNR) diet (*n* = 12), as previously described (Sohlstrom, Katsman et al. 1998, Elias, Maki et al. 2017). Time mating, pregnancy confirmation ultrasounds, and ultrasound-guided, transcutaneous, intra-placental nanoparticle treatment were performed as previously described (Wilson, Lampe et al. 2021). Briefly, nanoparticles were formed by complexing plasmids containing the *hIGF-1* gene under the control of the placental trophoblast specific promotor *CYP19A1* with a non-viral PHPMA_115_-b-PDMEAMA_115_ co-polymer (50 µg plasmid in a 200 µL injection volume) (Wilson, Lampe et al. 2021). At GD30-33, females underwent a nanoparticle injection of plasmid containing *hIGF-1* (Control + *hIGF-1* n = 4 or MNR + *hIGF-1* n = 7), or PBS sham injection (Control n = 7 or MNR n = 5), and were then sacrificed five days after injection (GD35-38). At time of sacrifice, major maternal and fetal organs were collected and weighed; fetal sex was also determined by visual inspection and confirmed using PCR as previously described (Depreux, Czech et al. 2018, Wilson, Stephens et al. 2021). Placentas were halved and either fixed 4% w/v paraformaldehyde (PFA) or the labyrinth and sub-placenta/decidua tissue separated, and snap-frozen in liquid nitrogen and stored at -80ºC.

### Placenta Quantitative PCR (qPCR)

Approximately 150-200 mg of snap frozen placenta labyrinth and sub-placenta/decidua tissue was lysed and homogenized in RLT-lysis buffer (*Qiagen*) using a tissue homogenizer. RNA was extracted and DNase treated using the RNeasy Midi Kit (*Qiagen*) following standard manufacturers protocols. The High-capacity cDNA Reverse Transcription kit (*Applied Biosystems*) was used to convert 1 µg RNA to cDNA, and qPCR reactions were set up and performed on the StepOne-Plus Real-Time PCR System (*Applied Biosystems*) as previously described (Wilson, Lampe et al. 2021). Commercially available KiCqStart (*Sigma*) primers were used. Relative mRNA expression was calculated using the comparative CT method with the StepOne Software v2.3 (*Applied Biosystems*).

### Immunohistochemistry (IHC)

Following fixation in 4% PFA, placenta tissue was processed and paraffin embedded following standard protocols. 5 µm thick tissue sections were obtained using a microtome (*Leica*), de-waxed and rehydrated as standard. Targeted antigen retrieval was performed by boiling sections in 1x IHC antigen retrieval solution (*Invitrogen*) for 15-20 mins, followed by a 20 min cooling period at room temperature. Incubation in 3% hydrogen peroxidase solution for 10 min was used to suppress endogenous peroxidase activity, and a protein block of 10% goat serum with 1% bovine serum albumin (BSA; *Jackson ImmunoResearch*) was the applied for 1 h at room temperature. Primary antibodies were diluted in 10% goat serum with 1% BSA and applied to sections overnight a 4ºC. Sections were then incubated with a biotinylated secondary antibody for 1 h at room temperature, amplified using the Vector ABC kit (*Vector*), and detected using DAB (*Vector*) Hematoxylin was used to counterstain nuclei. Details on antibodies and dilutions are supplied in Supplementary Table S2. Stained sections were visualized using the Axioscan scanning microscope (*Zeiss*), and representative images obtained using the Zen imaging software (*Zeiss*)

### Western Blot Analysis

Sub-placenta/decidua tissue was homogenized in ice-cold RIPA buffer containing protease and phosphatase inhibitors and protein concentrations determined using the Pierce™ Coomassie Blue Assay kit (*Thermo Scientific*) following standard protocol. 40 µg of protein was then mixed with Nupage SDS Loading Buffer (*Invitrogen*) and Reducing Agent (*Invitrogen*) and denatured by heating at 95ºC for 10 min. The lysates were then run on a 4-12% Tris-Bis precast gel (*Invitrogen*) following manufacturers protocols and transferred onto nitrocellulose membranes using the Bolt Mini Electrophoresis unit (*Invitrogen*). Both a Prestain Protein ladder (PageRuler, *Thermo Scientific*) and Biotinylated Protein ladder (*Cell Signaling*) were included on the gels. Membranes were placed into 5% skim-milk in Tris-buffered Saline containing Tween 20 (TBS-T) and incubated overnight at 4ºC. Primary antibodies (Supplemental Table S1) were then applied for 1-2 h at room temperature, the membranes washed 3 times in fresh TBS-T, and then further incubated with a HRP conjugated secondary (*Cell Signaling*) for 1 h at room temperature. During the secondary incubation, an anti-biotin antibody (*Cell Signaling*) was also applied to visualize the biotinylated protein ladder. Protein bands were visualized by chemiluminescence using SuperSignal West Femto Maximum Sensitivity Substrate (*Thermo Scientific*) on the Universal Imager (*Biorad*). Signal intensity of the protein bands were calculated using Image Lab software (version 6.1, *Biorad*) and normalized to β-actin.

### Statistical analysis

All statistical analyses were performed using SPSS Statistics 27 software. Generalized estimating equations were used to determine differences between diet and nanoparticle treatment, with maternal environment treated as a random effect and gestational age as a covariate. Fetal sex was also included as a main effect but removed if no statistical significance (P≤0.05) was determined. For statistically significant results, a Bonferroni post hoc analysis was performed. Results are reported as estimated marginal means ± standard error.

## Results

### *hIGF-1* nanoparticle treatment reduces pErk expression in control sub-placenta/decidua but increases pAkt expression in MNR sub-placenta/decidua

We have shown that sub-placenta/decidua expression of endogenous *Igf1* is not altered by either diet or *hIGF-1* nanoparticle treatment (Wilson, Lampe et al. 2021). Sub-placenta/decidua expression of *Igf1 Receptor* (*Igf-1R*), *IGF Binding Protein 3* (*IgfBP3*) and *Igf-2* were also not impacted by either diet or *hIGF-1* nanoparticle treatment (all P>0.05; Supplemental Figure 1).

IGF-1 binding to IGF-1R modifies gene expression and cellular function through the ERK/AKT signaling pathways. Both MNR and control + *hIGF-1* sub-placenta/decidua demonstrated reduced expression of pErk compared to Control (P=0.025 and P<0.001, respectively; Fig. 1A). When comparing MNR groups, there was no difference in pErk expression between MNR and MNR + *hIGF-1*. pAkt expression was not changed in Control + *hIGF-1* sub-placenta/decidua, however expression increased in the MNR + *hIGF-1* sub-placenta/decidua when compared to Control and MNR (P=0.038; Fig. 1B). Compared to Control, there was decreased protein expression of the mTORC1 component Raptor in Control + *hIGF-1* sub-placenta/decidua (P<0.01; Fig. 1C). However, there was no difference in Raptor expression in the MNR and MNR + *hIGF-1* when compared to Control (P>0.05; Fig. 1C), nor was there a change in the mTORC2 component Rictor (P>0.05; Fig. 1D). Protein expression of the mTOR inhibitor Deptor was increased in the Control + *hIGF-1* and MNR + *hIGF-1* sub-placenta/decidua compared to Control (P<0.05 for both, respectively; Fig. 1E).

**Figure 1.**
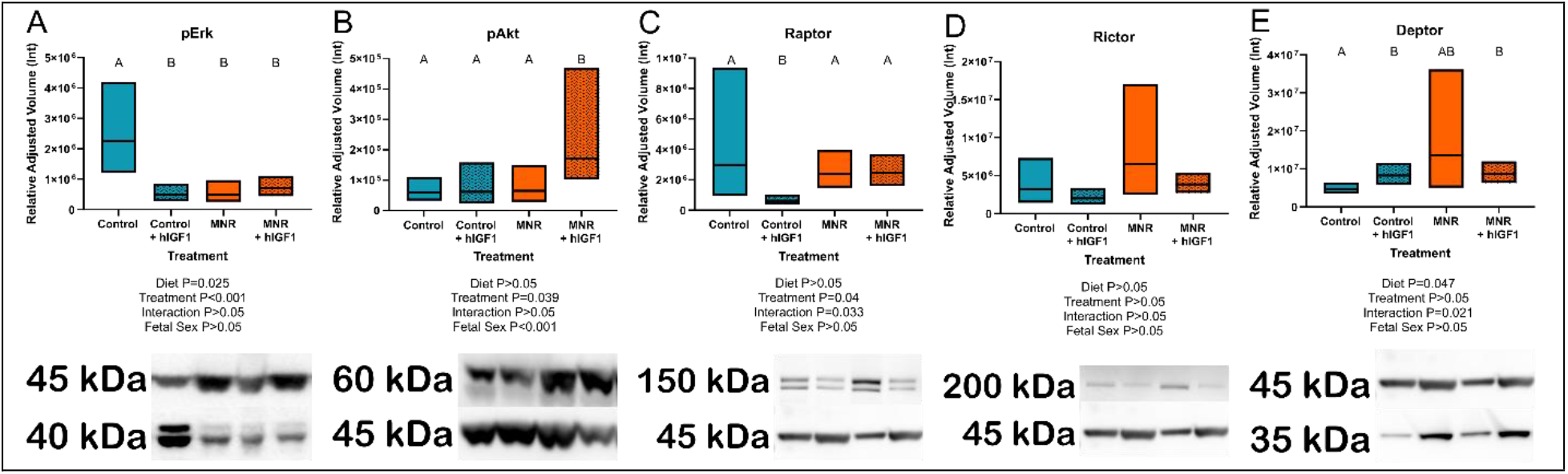
Effect of maternal nutrient restriction (MNR) diet and *hIGF-1* nanoparticle treatment on expression of Erk/Akt signaling proteins. **A**. Protein expression of phospho-Erk (pErk) was reduced in Control + *hIGF-1*, MNR and MNR + *hIGF-1* sub-placenta/decidua when compared to Control. **B**. In MNR + *hIGF-1* nanoparticle treatment there was increased expression of phospho-Akt (pAkt) compared to MNR, but there was no difference between Control + *hIGF-1* and Control sub-placenta/decidua. **C**. Compared to Control, there was decreased expression of Raptor in Control + *hIGF-1* nanoparticle treatment, but no difference with MNR or MNR + *hIGF-1* nanoparticle treatment **D**. There was no difference in the expression of Rictor across any groups. **D**. *hIGF-1* nanoparticle treatment increased expression of Deptor in both Control + *hIGF-1* and MNR + *hIGF-1* sub-placenta decidua when compared to Control. Representative western blots for each protein, left to right: Control, Control + *hIGF-1*, MNR, MNR + *hIGF-1*. n = 4 Control dams (8 sub-placenta/decidua), 4 Control + hIGF-1 dams (8 sub-placenta/decidua) 4 MNR dams (8 sub-placenta/decidua) and 4 MNR + hIGF-1 dams (8 sub-placenta/decidua). Data are estimated marginal means ± 95% confidence interval. P values calculated using generalized estimating equations with Bonferroni post hoc analysis. Different letters denote a significant difference of P≤0.05.

### *hIGF-1* nanoparticle treatment differentially regulates vascular growth factor mRNA expression in the placenta in a diet dependent manner

Angiogenic growth factor mRNA in the sub-placenta/decidua was differentially regulated by *hIGF-1* depending on maternal diet group (*VegfA*: P=0.049, *Pgf*: P<0.01, *PdgfB*: P<0.01, *VegfB*: P<0.001, *PdgfBR*: P=0.035; Fig. 2). *VegfA, Pgf* and *PdgfB* mRNA expression was decreased in Control + *hIGF-1* sub-placenta/decidua (P<0.05; Fig. 2A-C) compared to Control. However, there was no difference in *VegfA, Pgf* or *PdgfB* expression in the MNR + hIGF sub-placenta-decidua compared to MNR. Conversely, expression of *VegfB, VegfR1* and *PdgfBR* was increased in MNR + *hIGF-1* sub-placenta/decidua compared to MNR

**Figure 2.**
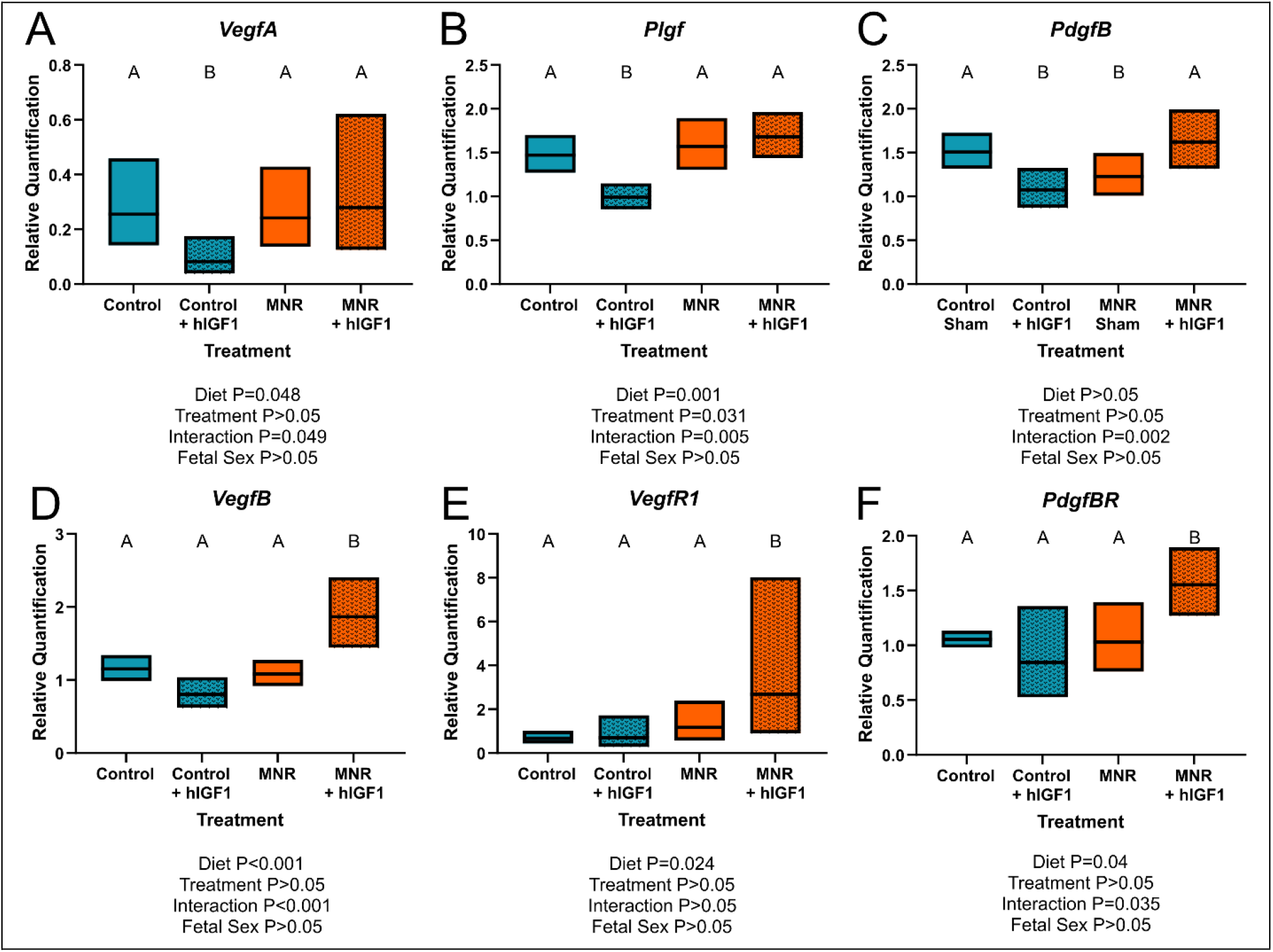
Effect of maternal nutrient restriction (MNR) diet and *hIGF-1* nanoparticle treatment on expression of angiogenic growth factors. Compared to Control, *hIGF-1* nanoparticle treatment reduced mRNA expression of *Vascular endothelial growth factor A* (*VegfA*; **A**) and *Placenta growth factor* (*Pgf;* **B**) in control + *hIGF-1* sub-placenta/decidua but was not different in MNR and MNR + *hIGF-1* nanoparticle treatment sub-placenta/decidua. **C**. Expression of *Platelet-derived growth factor B* (*PdgfB*) was decreased in Control + *hIGF-1* and MNR sub-placenta/decidua compared to Control, and increased in MNR + *hIGF-1* to normal. In MNR + *hIGF-1* nanoparticle treatment, there was increased expression of *VegfB* (**D**), *Vegf Receptor 1* (*VegfR1*; **E**) and *PdgfB Receptor* (*PdgfBR*; **F**) compared to all other groups. n = 6 Control dams (12 sub-placenta/decidua), 4 Control + *hIGF-1* dams (12 sub-placenta/decidua) 5 MNR dams (10 sub-placenta/decidua) and 7 MNR + *hIGF-1* dams (13 sub-placenta/decidua). Data are estimated marginal means ± 95% confidence interval. P values calculated using generalized estimating equations with Bonferroni post hoc analysis. Different letters denote a significant difference of P≤0.05.

### *hIGF-1* nanoparticle treatment normalizes expression of EMT factors in the MNR sub-placenta/decidua

IHC confirmed expression of Cldn5 in trophoblasts lining maternal spiral arteries within the sub-placenta/decidua (Fig. 3A), Abcb1 in the cytotrophoblasts within the cell columns (Fig. 3B), and Ocln in the trophoblast progenitor cells of the cell columns (Fig. 3C). Cytokeratin 7 was used to confirm expression of Cldn5, Abcb1 and Ocln was localized to trophoblasts (Fig. 3D & E). mRNA expression of *Cldn5* was reduced in the MNR sub-placenta/decidua (P=0.001; Fig. 4A), but was not regulated by *hIGF-1* treatment. In MNR + *hIGF-1* gene expression of *Abcb1* was increased when compared to all groups (P<0.05; Fig. 4B). Increased expression of *Ocln* and *Zo1* was evident in MNR compared to Control, but was normalized by *hIGF-1* treatment (P=0.01 & P=0.036; Fig. 4C & 4D, respectively). At the protein level, there was no difference in expression of Cldn5 with diet or *hIGF-1* nanoparticle treatment (Fig. 4E). *hIGF-1* nanoparticle treatment increased protein expression of Abcb1 in Control + *hIGF-1* and MNR sub-placenta/decidua compared to Control (P=0.047 & P=0.046, respectively; Fig. 4F). There was no difference in protein levels of Ocln with either diet or *hIGF-1* nanoparticle treatment.

**Figure 3.**
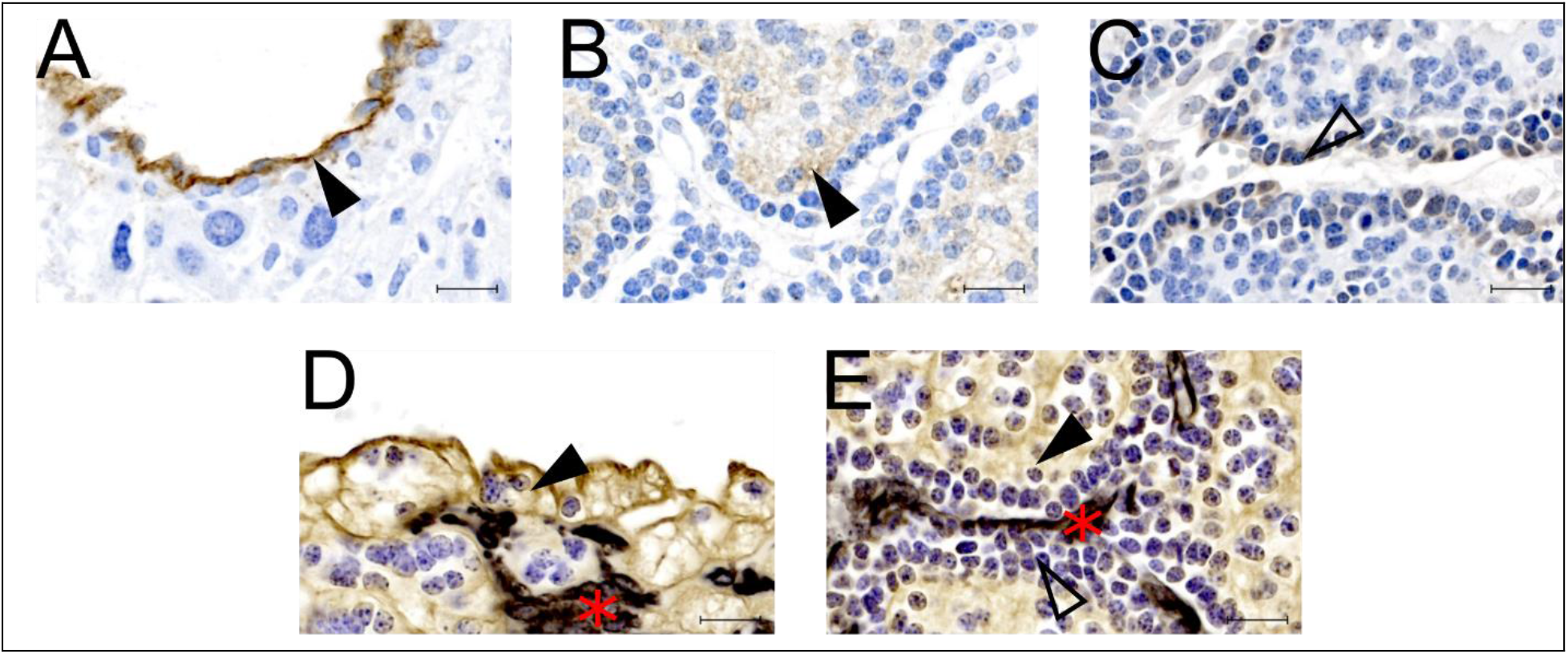
Protein localization of epithelial-mesenchymal transition (EMT) factors in the sub-placenta/decidua. **A**. Representative image of immunohistochemistry (IHC) staining of Claudin 5 in the sub-placenta/decidua trophoblasts (closed arrow head) that line maternal spiral arteries. **B**. Representative image of IHC staining of Abcb1 (P-glycoprotein) in the sub-placenta/decidua cytotrophoblasts (closed arrow head) of the cell column. **C**. Representative image of IHC staining of Occludin in the trophoblast progenitor cells in the cell columns (open arrow head). **D**. Representative image of double label IHC staining for Cytokeratin (brown; closed arrow head), indicating trophoblast cells, and Vimentin (black: red asterisk), indicating blood vessels, of a maternal spiral artery. **E**. Representative image of double label IHC staining for Cytokeratin (brown; closed arrow head), indicating cytotrophoblast cells, and Vimentin (black: red asterisk), indicating fetal blood vessels, of the sub-placenta cell column. Trophoblast progenitor cells in the cell column stained weakly for cytokeratin (open arrow head). Scale bar = 10 µm. n = 3-5 control placentas.

**Figure 4.**
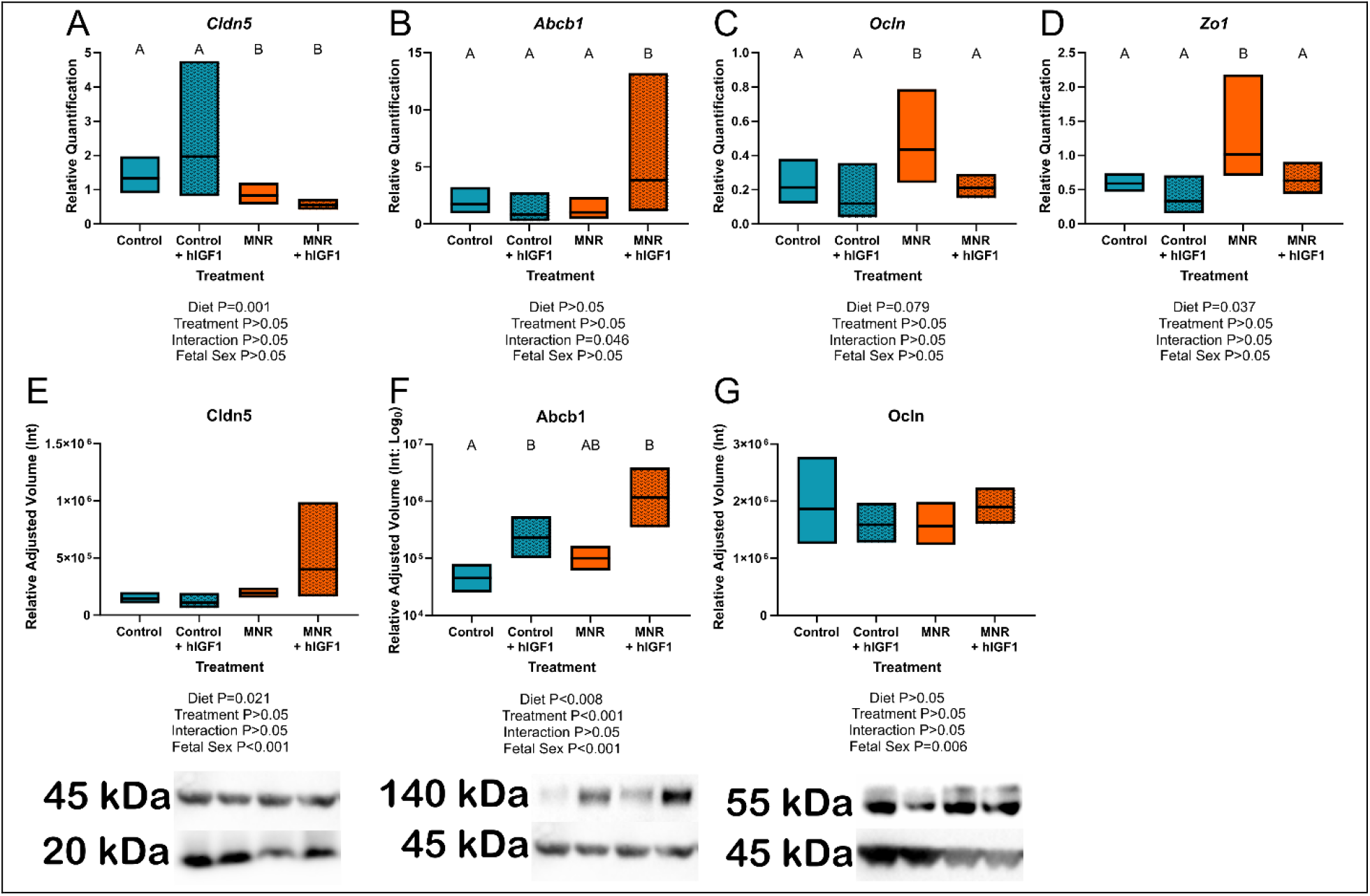
Effect of maternal nutrient restriction (MNR) diet and *hIGF-1* nanoparticle treatment on expression of epithelial-mesenchymal transition factors. **A**. There was decreased expression of *Claudin 5* (*Cldn5*) in MNR and MNR + *hIGF-1* sub-placenta/decidua compared to Control and Control + *hIGF-1* nanoparticle treatment **B**. There was increased expression of *P-glycoprotein* (*Abcb1*) in MNR + *hIGF-1* compared to Control, Control + *hIGF-1* and MNR. **C**. In MNR sub-placenta/decidua, there was increased expression of *Occludin* (*Ocln*; **C**) and *Tight junction protein 1* (*Zo1*; **D**) compared to Control, but expression was decreased in MNR + *hIGF-1* nanoparticle treatment to normal. **E**. Neither diet or *hIGF-1* nanoparticle treatment affected protein expression of Cldn5. **F**. Protein expression of Abcb1 was increased in Control + *hIGF-1* and MNR + *hIGF-1* nanoparticle compared to control. **G**. Neither diet or *hIGF-1* nanoparticle treatment affected protein expression of Ocln. Representative western blots for each protein, from left-right: Control, Control + *hIGF-1*, MNR, MNR + *hIGF1*. qPCR analysis: n = 6 Control dams (12 sub-placenta/decidua), 4 Control + *hIGF-1* dams (12 sub-placenta/decidua) 5 MNR dams (10 sub-placenta/decidua) and 7 MNR + *hIGF-1* dams (13 sub-placenta/decidua). Western blot: n = 4 Control dams (8 sub-placenta/decidua), 4 Control + *hIGF-1* dams (8 sub-placenta/decidua) 4 MNR dams (8 sub-placenta/decidua) and 4 MNR + *hIGF-1* dams (8 sub-placenta/decidua). Data are estimated marginal means ± 95% confidence interval. P values calculated using generalized estimating equations with Bonferroni post hoc analysis. Different letters denote a significant difference of P≤0.05.

## Discussion

The etiology of fetal growth restriction (FGR) is often associated with aberrant extravillous trophoblast (EVT) invasion which limits maternal blood flow and thus nutrient/oxygen availability (Abbas, Turco et al. 2020). The aim of the present study was to determine the effects of maternal nutrient restriction (MNR) on sub-placenta trophoblast signaling, and whether increasing *hIGF-1* expression, with nanoparticle-mediated gene therapy impacted ERK/AKT signaling pathways and expression of factors involved in EMT and vascular remodeling. We show that MNR results in reduced levels of pErk in the sub-placenta/decidua and was associated with reduced gene expression of *PdgfB*, and increased expression of *Ocln* and *Zo1*. In MNR placentas treated with *hIGF-1* nanoparticle, expression of *PdgfB, Ocln* and *Zo1* was normalized, whilst expression of *VegfB, Vegf-R1* and *PdgfB-R* was increased.

The ERK/AKT signaling pathways are complex, involving multiple protein families, and can be activated or inactivated through different phosphorylation mechanisms (Bhaskar and Hay 2007, Fremin, Saba-El-Leil et al. 2015). In the present study, MNR reduced the expression of pErk in the sub-placenta/decidua compared to Control, but had no effect on pAkt expression. Homozygous knockout of ERK isoform 1 (*Erk1*^*-/-*^), results in little disruption to embryonic development, with fetuses born viable (Pages, Guerin et al. 1999), however, homozygous knockout of isoform 2 (*Erk2*^*-/-*^) results in embryonic lethality at E6.5 despite expression of ERK1 in the embryos (Hatano, Mori et al. 2003). Loss of a single *Erk2* allele results in decreased placental weight, which is further exacerbated with disruption to *Erk1* (Fremin, Saba-El-Leil et al. 2015), and emphasizes the role of placental ERK1/2 activity during gestation. pErk expression remained reduced compared to control in the MNR + *hIGF-1* sub-placenta/decidua, however there was increased expression of pAkt with *hIGF-1* nanoparticle treatment, indicating the ability to manipulate ERK/AKT signaling in the dysfunctional placenta.

In normal human pregnancies, EVT invasion of the maternal vasculature result in remodeling of the maternal spiral arteries into low resistance vessels which allows for delivery of high volumes of nutrient and oxygen rich maternal blood to enter the placenta and support adequate fetal growth (Boyd and Hamilton 1970). However, in pregnancies complicated by FGR, insufficient spiral artery remodeling is often a common feature of placental pathology (Burton, Woods et al. 2009). We have previously shown that *hIGF-1* nanoparticle treatment of the placenta increases *hIGF-1* in trophoblasts residing in the sub-placenta/decidua (Wilson, Lampe et al. 2021). These trophoblasts in the guinea pig placenta are responsible for communication with the maternal environment and remodeling of the maternal vasculature. Utero-Placental blood flow in the MNR guinea pig placenta remains to be determined, however, MNR decreased placental blood flow has been shown in rat MNR pregnancies (Ahokas, Anderson et al. 1983). In MNR + *hIGF-1* sub-placenta/decidua, there was increased expression of pAkt which was associated with increased expression of *VegfB, Vegf-R1 PdgfB* and *PdgfB-R*. VEGF and PDGFB are produced by the placenta and secreted into maternal circulation where they aid maternal adaptations to pregnancy like inducing vasodilation of resistance vessels (Chen, Liu et al. 2015, Cuffe, Holland et al. 2017). VEGF in particular, has been shown to act as a vasodilator for myometrial resistance arteries (Szukiewicz, Szewczyk et al. 2005) and is decreased in maternal serum of FGR pregnancies (Nadar, Karalis et al. 2005). Furthermore, VEGF and PDGFB play an important role in regulating the uterine immune response involved in regulating trophoblast invasion (Kalkunte, Mselle et al. 2009, Wang, Nian et al. 2021). Increased expression of *VegfB, PdgfB* and their receptors did not result in noticeable changes to the gross morphology of the sub-placenta/decidua examined by 2D histology (Wilson, Lampe et al. 2022). This may be due to the short time period between *hIGF-1* nanoparticle treatment and sample collection, and with longer treatment may result in noticeable changes to trophoblast invasion and thus maternal spiral artery remodeling.

Tight junction and adhesion proteins, including claudins, occludins, and ZO-1, are critical components in determining epithelial cell polarity and are repressed in trophoblasts undergoing EMT prior to trophoblast invasion (Marzioni, Banita et al. 2001, Ikenouchi, Matsuda et al. 2003). In the guinea pig sub-placenta/decidua, Cldn5 was expressed specifically in trophoblasts lining the maternal spiral arteries, Abcb1 was expressed in the trophoblasts of the sub-placenta cell columns, and Ocln to the trophoblast progenitor cells. The cell columns of the guinea pig sub-placenta contain the progenitor trophoblast that become invasive trophoblast responsible for maternal vascular remodeling (Verkeste, Slangen et al. 1998). Under MNR conditions we demonstrated reduced *Cldn5* expression, and increased *Ocln* and *Zo1* expression in the sub-placenta/decidua suggesting strong cell adhesion and tight junctions which would not be conducive to EMT and invasion. However, *hIGF-1* expression in the MNR sub-placenta/decidua resulted in normalization of *Ocln* and *Zo1* levels to that of Control, and expression of *Abcb1* was increased compared to MNR at the transcript but not protein level. This may be due to the short time period of treatment or the use of whole sub-placenta/decidua samples, although single cell analysis is beyond the scope of this manuscript.

Currently, more than 50% of FGR cases remain undiagnosed prior to birth due, in part, to a lack of definitive ways to predict and detect FGR (Sun, Groom et al. 2020). The placenta is an ideal target for treatment of FGR as it contributes significantly to the etiology of FGR, and is discarded after birth, thus mitigating any potential long-term consequences for maternal health. We have previously shown that treatment of the placenta with *hIGF-1* nanoparticle does not result in maternal or fetal toxicity (Wilson, Lampe et al. 2022). Furthermore, in the present study we show that treatment of control placentas with the *hIGF-1* nanoparticle results in reduced expression of pErk, *VegfA* and *Pgf*, and increased expression of the mTOR inhibitor Deptor indicating the ability of the sub-placenta/decidua under Control conditions to maintain homeostasis when excess IGF-1 is present.

In conclusion, we have shown that under maternal nutrient restrictions (MNR), *hIGF-1* nanoparticle treatment triggers increased expression of growth factors and normalization of EMT factors, which are likely to positively influence the development, structure and function of the maternal-fetal interface. However, under control conditions, the response of the placenta is to reduce AKT/mTOR signaling and growth factor expression to maintain homeostasis.

## Acknowledgments

We would like to thank Drs Craig Duvall and Mukesh Gupta for providing the co-polymer, and Kristin Lampe for her assistance with the animal experiments.

## Contributions

BD performed experiments, analyzed data and wrote manuscript. HNJ obtained funding, conceived the study and edited manuscript. RLW conceived the study, performed experiments, analyzed data and wrote manuscript. All authors approve final version of manuscript.

## Ethics approval

Animal care and usage was approved by the Institutional Animal Care and Use Committees at Cincinnati Children’s Hospital and Medical Center (Protocol number 2017-0065).

## Competing Interests

The authors have declared that no competing interest exists

## Data availability

All data needed to evaluate the conclusions in the paper are present in the paper and/or the Supplementary Materials.

## Funding

This study was funded by Eunice Kennedy Shriver National Institute of Child Health and Human Development (NICHD) award R01HD090657 (HNJ).

## Notes

### Competing Interest Statement

The authors have declared no competing interest.

