## Supplementary Material for "Placental treatment with *insulin-like growth factor 1* via nanoparticle differentially impacts vascular remodeling factors in guinea pig sub-placenta/decidua"

**Supplemental Material**

| **Supplemental Table S1** Primary antibodies and concentrations used in Western Blot and Immunohistochemistry (IHC) | | |
| --- | --- | --- |
| **Protein** | **Manufacturer** | **Concentration** |
| Phospho-Erk1/2 | Cell Signaling *4370* | 1:500 |
| Phospho-Akt1 (S473 & T308) | Abcam *Ab66138* | 1:1000 |
| Raptor | Cell Signaling *2280S* | 1:1000 |
| Rictor | Cell Signaling *2114S* | 1:1000 |
| Deptor | LSBio *LS-C187268* | 1:1000 |
| Claudin 5 | Invitrogen *35-2500* | 1:500 (Western) & 1:50 (IHC) |
| Abcb1/P-glycoprotein | LSBio *LS-B5171* | 1:1000 (Western) & 1:25 (IHC) |
| Occludin | Invitrogen *40-4700* | 1:500 (Western) & 1:50 (IHC) |

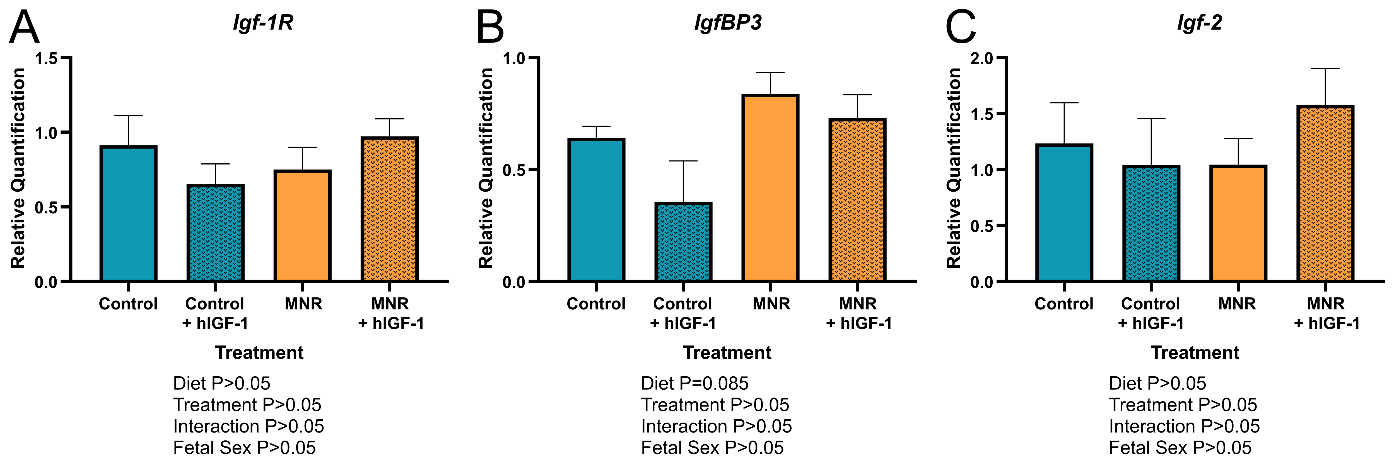

**Supplemental Figure 1. Effect of maternal nutrient restriction (MNR) diet and *hIGF-1* nanoparticle treatment on expression of insulin-like 1 (Igf-1) growth factor signaling member.** There was no difference in the expression of *Igf1* *Receptor* (*Igf-1R*; **A**), *IGF Binding Protein 3* (*IgfBP3*; **B**) and *Igf2* (**C**) with either diet or *hIGF-1* nanoparticle treatment. Data are estimated marginal mean ± standard error. Control dams (12 sub-placenta/decidua), 4 Control + *hIGF-1* dams (12 sub-placenta/decidua) 5 MNR dams (10 sub-placenta/decidua) and 7 MNR + *hIGF-1* dams (13 sub-placenta/decidua). Data are estimated marginal means ± 95% confidence interval. P values calculated using generalized estimating equations with Bonferroni post hoc analysis.
